# Robust Algorithms for Capturing Population Dynamics and Transport in Oceanic Variables along Drifter Trajectories using Linear Dynamical Systems with Latent Variables

**DOI:** 10.1101/438036

**Authors:** Yan Yan, Tony Jebara, Ryan Abernathey, Joaquim Goes, Helga Gomes

## Abstract

The blooms of *Noctiluca* in the Gulf of Oman and the Arabian Sea have been intensifying in recent years posing a threat to regional fisheries and the long-term health of an ecosystem supporting a coastal population of nearly 120 million people. We present the results of a microscopic data analysis to investigate the onset and patterns of the *Noctiluca* (mixotrophic dinoflagellate *Noctiluca scintillans*) blooms, which form annually during the winter monsoon in the Gulf of Oman and in the Arabian Sea. Our approach combines methods in physical and biological oceanography with machine learning techniques. In particular, we present a robust algorithm, the variable-length Linear Dynamic Systems (**vLDS**) model, that extracts the causal factors and latent dynamics at the microscopic population-level along each individual drifter trajectory, and demonstrate its effectiveness by using it to test and confirm previously benchmarked macroscopic scientific hypotheses. The test results provide microscopic statistical evidence to support and recheck the macroscopic physical and biological Oceanography hypotheses on the *Noctiluca* blooms; it also helps identify complementary microscopic dynamics that might not be visible or discoverable at the macroscopic scale. The vLDS model also exhibits a generalization capability (inherited from a machine learning methodology) to investigate important causal factors and hidden dynamics associated with ocean biogeochemical processes and phenomena at the population-level.

## Introduction

### Background

Recent advances in Data Science and Machine Learning have produced great successes in a variety of data-driven modeling for interdisciplinary scientific problems concerning complex natural phenomena, in Marine Ecology [1–6], Climatology [7], Oceanography [8–11], Geoscience [12], Computer Vision [13–15], Social Science [16], Computational Neuroscience [17–20], Speech and Language Processing [21–23], and Environmental Health Science [24]. Here we use these techniques to investigate the onset and patterns of the *Noctiluca* winter monsoon blooms, which form annually and with predictable regularity in the Gulf of Oman and in the Arabian Sea. Our approach relies on a combination of satellite and drifter derived oceanographic data and machine learning techniques. In particular, we obtain a robust model (the vLDS mode, variable-length Linear Dynamic System Model) that is capable of identifying the causal factors and dynamics at the microscopic population-level along each individual drifter trajectory. Furthermore, we assess its effectiveness via testing and confirm previously benchmarked scientific hypotheses. Rigorously statistical, the vLDS model is a powerful tool that helps us: 1) discover microscopic causal relationships in a high-dimensional dataset and 2) identify complementary microscopic dynamics that might not be discoverable at the macroscopic scale or accessible in controlled laboratory experiments.

The significance of this research is that these blooms of *Noctiluca* have been intensifying in recent years posing a threat to regional fisheries and the long-term health of an ecosystem supporting a coastal population of nearly 120 million people [25–28]. When seen from space, the *Noctiluca* blooms appear as large drifting swirls and filaments on the surface of the sea (Fig 1A). Traditionally, photosynthetic diatoms supported the Arabian Sea food chain; zooplankton preyed primarily on diatoms, which were in turn grazed by fish. Since early 2000s, the ecosystem of the Arabian Sea appears to have changed [27], with large and widespread blooms of *Noctiluca* superseding diatoms which dominated winter monsoon blooms. Gomes et al. [26] were able to discover that the annual outbreaks of *Noctiluca* were linked to the up shoaling of hypoxic waters from depth into the euphotic zone. Within a decade and half, *Noctiluca* blooms have virtually replaced diatoms at the base of the food chain, marking what appears to be an unprecedented ecosystem shift.

In previous studies, Gomes et al. [25–26], Goes et al. [28] relied on satellite observations and in-situ data sampling and biologically controlled experiments on board of research vessels, to describe the underlying dynamics governing the transport, the growth and decay of the *Noctiluca* blooms in the Arabian Sea region. It has been demonstrated that *Noctiluca* is capable of migrating up and down in the water column depending on conditions at the surface. Like all dinoflagellates [29–32], a stable water column is essential for the growth and proliferation of *Noctiluca* as large surface blooms. Since it is a mixotroph, it can meet its metabolic requirements via feeding on external prey or through photosynthesis by thousands of green “endosymbionts,” living within its central symbiosome (Fig 1B). This flexibility gives it an edge on diatoms, which survive on sunlight alone. When feeding on an external source of organic matter Noctiluca tends to accumulate significant amounts of ammonia [33] and lipids [27], making them highly buoyant, allowing them to accumulate at the sea surface for prolonged periods of time. When present at the surface *Noctiluca* can be easily transported by physical oceanographic processes, where there appear in satellite ocean color in association with micro and meso-scale eddies, filaments and streamers. Over the past decade in particular, *Noctiluca* blooms have become more intense and widespread. The blooms of 2015 and 2017 were the largest on record occupying an area almost thrice the size of the State of Texas.

Summarizing this complex biogeochemical process, recent research [26–27] provided indications that the biological or environmental trigger for the *Noctiluca* blooms were nutrient rich, low-oxygen waters, brought to the surface by an annually recurring mesoscale cyclonic eddies. The eastward propagation of these eddies and filaments associated with them appear to be responsible for the gradual spread of the bloom from their inception in the Sea of Oman in Dec. towards the east by January until mid-March (Fig 1C). The underlying forces responsible for the dispersal of the winter-time *Noctiluca* blooms are multiple. While *Noctiluca’s* unique physiological properties play an important role in maintaining the blooms at the surface and the overall integrity of the bloom, its dispersal is primarily due to ocean currents and the directions in which they move and mix with other water masses [34–39]. This mixing at meso-and sub-mesoscale occurs and changes over a seasonal basis, due to the interactions between the ocean and atmosphere that are driven by the monsoonal wind forcing [40–41].

**Fig 1.**
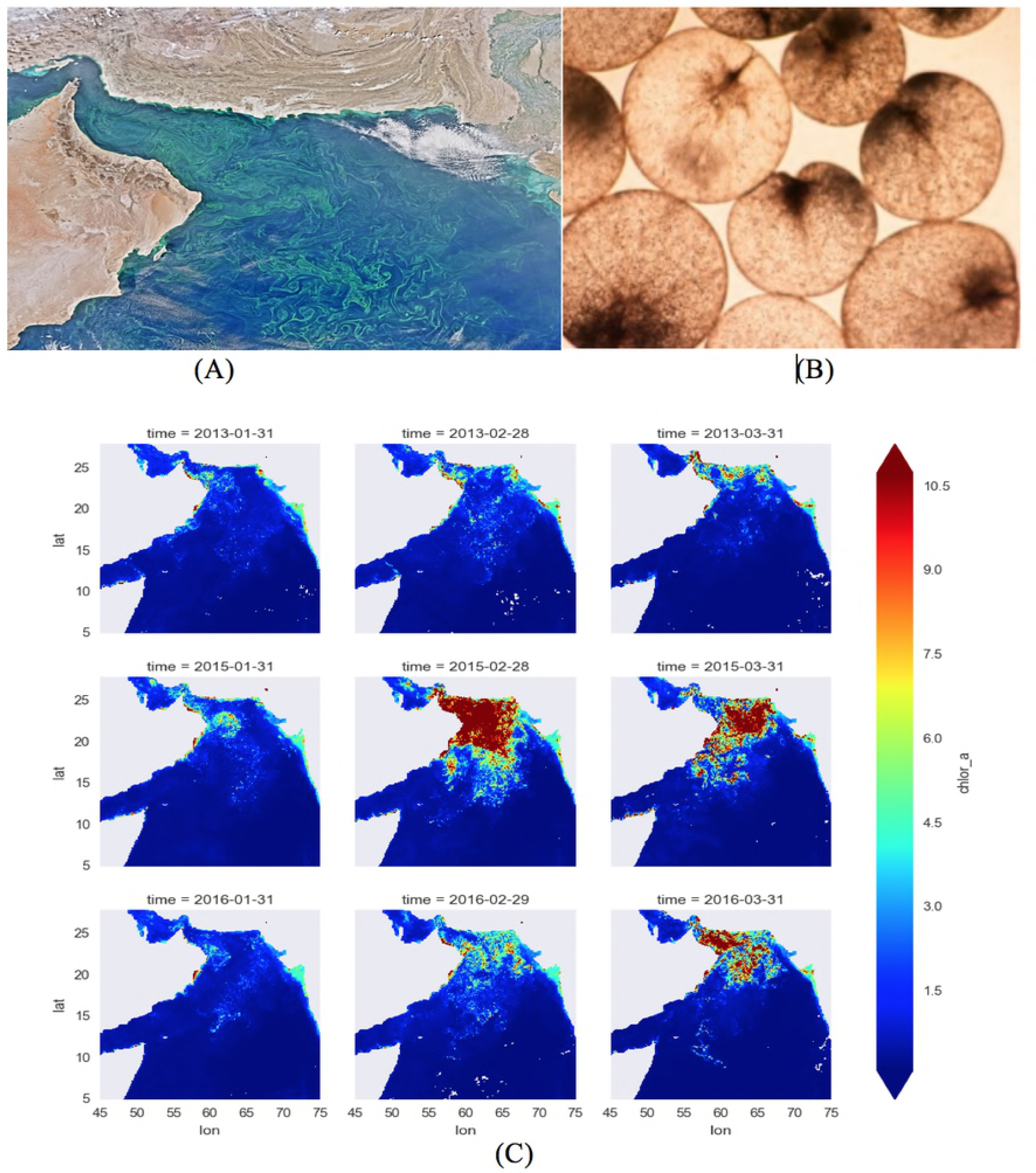
**(A)** Satellites images displaying Noctiluca blooms as large swirls on the surface of the Arabian Sea [42]. **(B)** Noctiluca scintillans with a flick of its tail-like flagellum drawing energy from the millions of green algae, or “endosymbionts”, captured inside its transparent cell walls. **(C)** Satellite image for the monthly data of the chlorophyll *a* concentration in the Arabian Sea during year 2013, 2015, and 2016.

Based on our previous research [26] and given *Noctiluca’s* rise in the Arabian Sea to an oxygen-starved hypoxia zone of more than 200 thousand square miles, we first (1) hypothesized and tested that nutrient-enrichment of the surface waters are increasing productivity and expanding the hypoxia zone as oxygen gets depleted. The potential sources of nutrients are multiple [25–26]. They include the domestic and industrial fallout from countries bordering the Arabian Sea, winter convective mixing during Northeast Monsoon (NEM, Nov-Jan) which helps bring up nutrients from the mixed layer into the euphotic layer of the Arabian Sea, and the atmospheric deposition measured by satellite as the aerosol optical thickness at wavelength (865 nm). As *Noctiluca* grows, it uses up oxygen in the water when feeding on other phytoplankton. Grazing on other phytoplankton and declining oxygen levels allows *Noctiluca* to outcompete diatoms during winter. We then (2) hypothesized and tested that *Noctiluca* grew faster in light than in dark on the sea surface and in the sea water, thanks to its sun-loving endosymbiotic algae, which are thought to have survived 1.3 billion years since being on an oxygen-scarce Earth.

### Goal and outline

To understand the microscopic impact of the biological and physical oceanographic factors on *Noctiluca* blooms along drifter trajectories, we have collected, combined, and preprocessed data from both the Ocean and Satellite datasets. The trajectory of each drifter is recovered by its spatio-temporal information. The biological and physical oceanographic profiles associated with the spatio-temporal coordinates of each drifter is then utilized to discern the the behavior and movement of the *Noctiluca* blooms along the trajectory of the particular drifter. This behavior and movement is statistically learned or transformed by the vLDS model. Since the drifters have different launch time and longevity, the time series representing individual drifter trajectories have different lengths. It is our goal in this paper to present the variable-length Linear Dynamic Systems (vLDS) model that is tailored to this particular data structure, to learn, summarize, and recover the latent dynamics for all drifter trajectories, and to generate predictive dynamics that match most closely the observed data along drifter trajectories.

The effectiveness of the vLDS model is demonstrated by testing our previous two hypotheses in the “Background” Section from the microscopic level and on both the physical and biological dynamics of the *Noctiluca* blooms. Moreover, *a priori* knowledge of the unique trait of *Noctiluca* is used in this study. The satellite based datasets that most closely align with the growth of *Noctiluca* blooms are colored dissolved organic material (*CDOM*) content of sea water, photosynthetically available radiation and diffuse attenuation coefficient at 490 nm using the Lee algorithm (light on and under the sea surface) and aerosol optical thickness over water (dust over the sea), and 4-micron nighttime sea surface temperature (*SST4*). The physical dispersion of the drifters or particles on the ocean surface is described by its spatio-temporal coordinates, eastward and northward velocity components, speed, and distance to the coast. The analysis of the phytoplankton blooms in [25–26, 40–41] is on the macroscopic scale of space and time, namely, the data is aggregated or pooled across spatio-temporal dimensions. In this study, the surface velocity drifter dataset [43] and satellite image dataset [44–47] are not aggregated or pooled across spatio-temporal dimensions. With the data structure of individual drifter trajectories kept intact, the vLDS model tests and works under our previous hypotheses on the dynamics of the *Noctiluca* blooms from the microscopic scale of space and time in the Arabian Sea region. More specifically, the vLDS model summarizes and captures the statistical structure of all drifter trajectories, by maximizing the log-likelihood of the joint distribution of the observations and their latent state variables and allowing each individual drifter trajectory generated by the model to vary as smooth functions in the latent dynamical state. By comparing the vLDS model predictions directly with the observed ocean profiles along the drifter trajectories, we can easily visualize its predictive performance and interpret the underlying dynamics of the *Noctiluca* blooms at the microscopic scale, as discussed in the “Discussion & conclusion” Section.

The benefit of vLDS is threefold. First, it provides statistical evidence in a direct and zoomed-in manner to support and recheck the macroscopic physical and biological oceanographic observations and the inherent physiological behavior of *Noctiluca* blooms, by recovering the latent dynamics that governs the probabilistic distribution of the *Noctiluca* concentration in space and time and by comparing the model predictions with the observed ocean profiles. Our study shows that most of the macroscopic hypotheses on the *Noctiluca* blooms can be statistically validated by the vLDS model. Second and more importantly, it helps us to identify complementary microscopic dynamics that might not be visible or discoverable at the macroscopic scale. The vLDS predictive plots in “Discussion & Conclusion” Section provide statistical evidence that the atmospheric deposition measured by the quantities *T865* aerosol optical thickness at wavelength 865 nm does not have much impact on the underlying dynamics that are driving the *Noctiluca* growth that is measured by the chlorophyll *a* concentration (*Chl a*) at the time scale of two days. Third, it provides a generalizable machine learning methodology to probe important microscopic causal factors and hidden dynamics for the ocean biogeochemical processes at the population-level along individual drifter trajectories. These scientific findings in the microscopic scale of the population can lead to critical hypothesis and even conclusions at the macroscopic scale of the pooled data. We note that this microscopic approach has been successfully applied in other areas, such as computational neuroscience [19–20] and sound tracking [22].

The major assumption of the **vLDS** model is that all the drifter trajectories are governed by the same set of biological and physical oceanographic principles or dynamics, in other words, all drifter trajectories in the vLDS model share the same model parameters. This assumption is both natural and physical. Those model parameters can then be learned jointly from the data, even if each drifter trajectory is evolving independently under the shared parameters to capture the autocorrelation and the latent dynamics in the full spatio-temporal, biological, and physical feature space. The model’s ability to capturing the underlying physical and biological dynamics of the *Noctiluca* blooms is quantified by the R-squared (*R*^2^) of the vLDS model fitted on the cross-validation dataset and predicted on both the cross-validation and heldout data. After the vLDS model parameters are learned from the data, we observed that the the vLDS model, recovers all the spatio-temporal variables well along drifter trajectories, as well as most of the biological factors, namely, the *CDOM*, light on and under the sea surface, *SST4*, except for the atmospheric deposition *T865*.

These test results support the claim that the assumption on the vLDS model is natural. Moreover, the recovered, summarized, and predicted latent dynamics (rather than the observed time series) has shown a close relationship among *Noctiluca* blooms, physical dispersal, and biological environments. The highly correlated relationships between the *Chl a* and *CDOM*, and between the *Chl a* and diffuse attenuation coefficient at 490 nm using the Lee algorithm (*KD490*, light under the sea surface) are close to linear. The tightly correlated relationships between the *Chl a* and photosynthetically available radiation (*PAR*, light on the sea surface) and between the *Chl a* and *SST4* are nonlinear. The aerosol optical thickness at wavelength 865 nm (*T865*) does not have much impact on the vLDS recovered dynamics or the *Chl a*. This corresponds to our hypotheses in the “Background” Section, and demonstrates that the nutrient and light are two important positive factors for the *Noctiluca* blooms and that the atmospheric deposition does not contribute much to the *Noctiluca*’s growth. Our two hypothesis are thus tested and confirmed by these test results.

## Materials and methods

### Data collection

The Arabian Sea (coordinate range ~5 to 28° N, 45 to 75° E) is predominantly located in the tropics (Fig 1A), and it has one of the most energetic current systems driven by the seasonally reversing monsoons. Due to the temperature gradient across the land mass and ocean, the winter monsoon from November to February is cold and dry and is blowing from the Indian subcontinent. Moreover, the winter monsoon cools the surface of the ocean and induce convective mixing that brings nutrients from depth to the surface. Nutrients and low oxygen waters brought to the surface by winter convective mixing during the winter monsoon foster large *Noctiluca* blooms of the Arabian Sea [26].

Datasets on *Chl a*, from the ocean color satellites Sea-WiFS, MODIS-Aqua (NASA), VIIRS (NOAA) and MERIS and OLCI-A (ESA), were used for studying the distribution of *Noctiluca* blooms during winter. In practice, the Ocean color datasets have missing values at certain locations due to the limitation of the satellite coverage (and also to the presence of clouds). The dark spots are regions with missing values (Fig 2). For the purpose of our study, we used merged products from both NASA and the GlobColour Project [44–47]. Ocean color satellites can provide remote sensing reflectance values for different wavebands. These wavebands are used in empirical and semi-analytical algorithms to convert remote sensing reflectance to chlorophyll *a* concentration. Pre-processed *Chl a* data products were used to explore a time series of snapshots of chlorophyll *a* concentration on a lattice of latitude and longitude coordinates. In the next sections, we provide detailed descriptions of the data aggregation and preprocessing steps.

**Fig 2.**
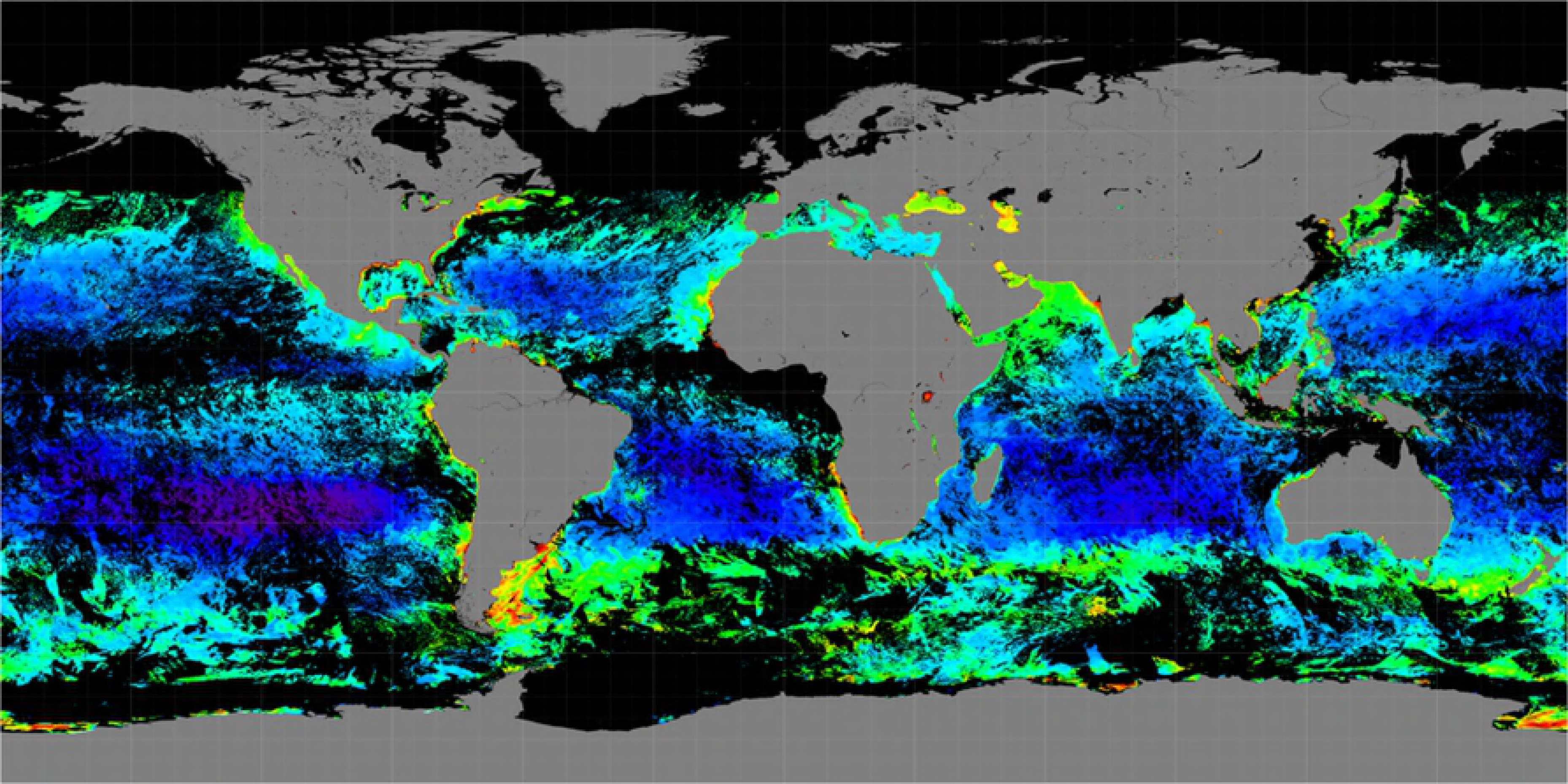
Level-3 data for the chlorophyll *a* concentration from Dec. 27 to Dec. 31, 2015 [42]. The dark regions indicate missing values.

The temporal evolution of the satellite images reflects both physical and biological dynamics. To impose the structures of physical drivers (advection) onto the data sample, we utilized the drifter array data from the NOAA’s Global Drifter Program (GDP). These freely drifting buoys provide information about the upper ocean currents that are responsible for the advection of the planktonic particles [48]. The Lagrangian trajectory of each float can be retrieved from the database as a time series of variables representing the location, velocity field, and sea surface temperature. We note that each float has a typical lifetime of a couple years and has different launch times. Therefore, the drifter dataset is highly heterogeneous in both time and space. In Fig 3, we display the temperature measurements of all the drifters in the Arabian Sea. It is known that the Lagrangian drifter trajectories are highly chaotic [49–51], and the prediction of particle trajectories has been a challenging research task [52]. There have been recent research results on using Kalman filter and data assimilation with various specific physical model, namely, the Gauss–Markov Lagrangian particle model [8], the Eulerian velocity field [9, 11], and the upper ocean horizontal momentum balance model from Ekman dynamics [10], to track the position and velocity of the floats. For comparison in our study, we are introducing and imposing statistical structure on the latent state variables to capture the joint dynamics among the *Chl a*, the spatio-temporal information of the floats, namely, the latitude, longitude, velocity, speed and distance to the coast, and the biological predictors, such as *CDOM, KD490, T865, PAR*, and *SST4*. The predictive plots generated by the vLDS model in our study (as displayed in Section “Discussion & Conclusion”) reveal the relationships among the physical, biological ocean profiles and the *Noctiluca* blooms inside a collection of chaotic drifter trajectories in the Arabian Sea region from 2002 to 2017.

**Fig 3.**
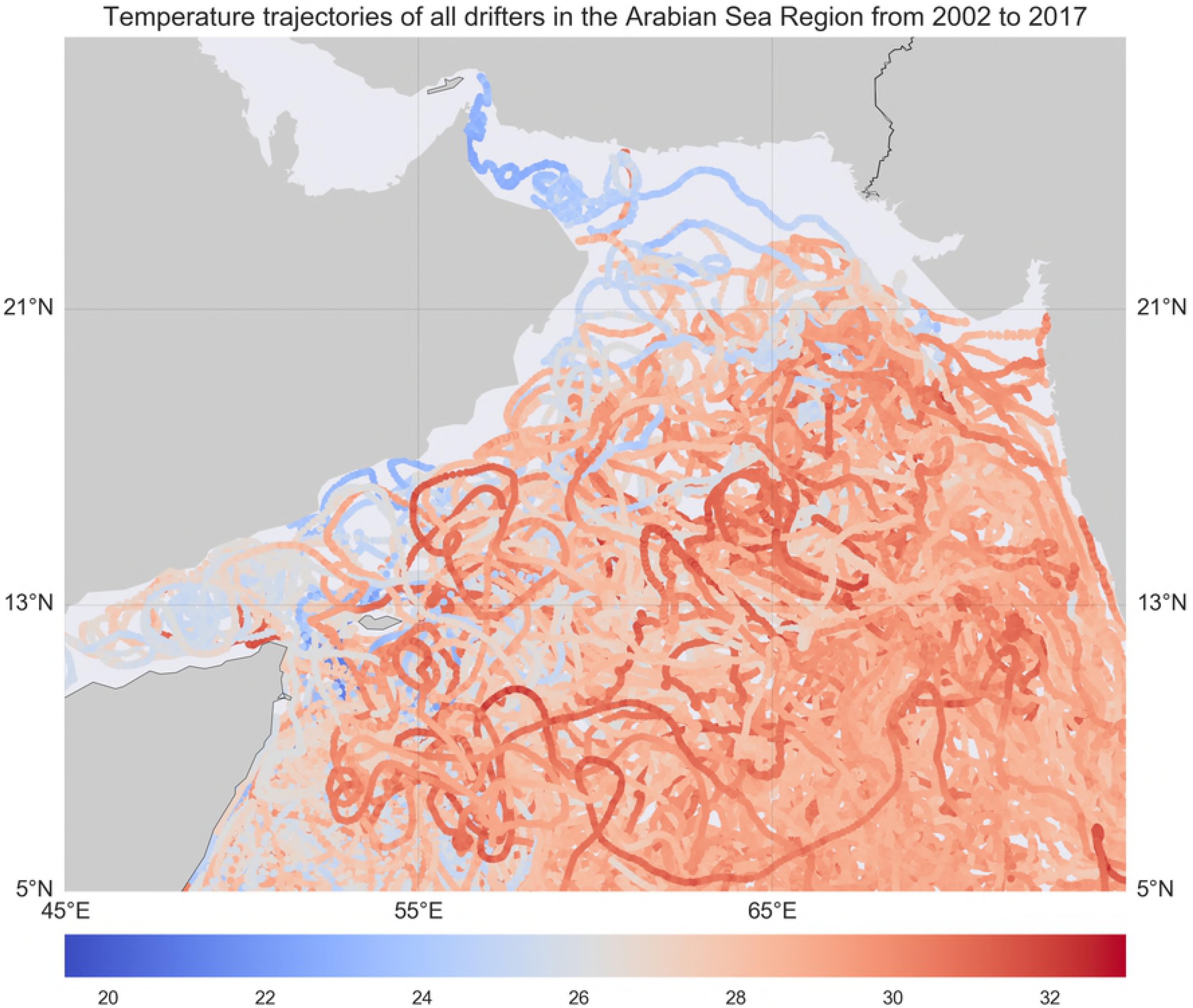
Temperature trajectories of all drifters in the Arabian Sea Region from 2002 to 2017.

### Combining multiple dataset

We merged the satellite data with the buoy data to generate a Lagrangian dataset. It is a collection of multivariate time series for each drifter with a unique id to combine the information from the satellites, namely, *Chl a, CDOM, KD490, T865, PAR*, and *SST4*, and the data associated with the drifters including *id, time*, latitude, longitude, velocity components, speed, and distance to the coast. Fourteen features were selected in our experiments (Table 1). The drifter *id* and *time* are mainly used for ordering and grouping data in the vLDS model. The twelve (12) remaining factors represent the physical and biological variables that related to the evolution of *Noctiluca* blooms [25–26]. Since the timescale of the phytoplankton reproduction is of the order of a few days [53], we carried out a resampling process to match the frequency of both datasets to the same level. At the same frequency, we interpolated the variables from the satellite dataset onto the specific spatial and temporal points of the Lagrangian drifter dataset, to add more information for each observation in the drifter dataset. This process builds up a multivariate time series for each drifter id with all the biological and physical information embedded on the drifter trajectory. Further, this interpolation process is repeated for all other features from the satellites.

**Table 1.**
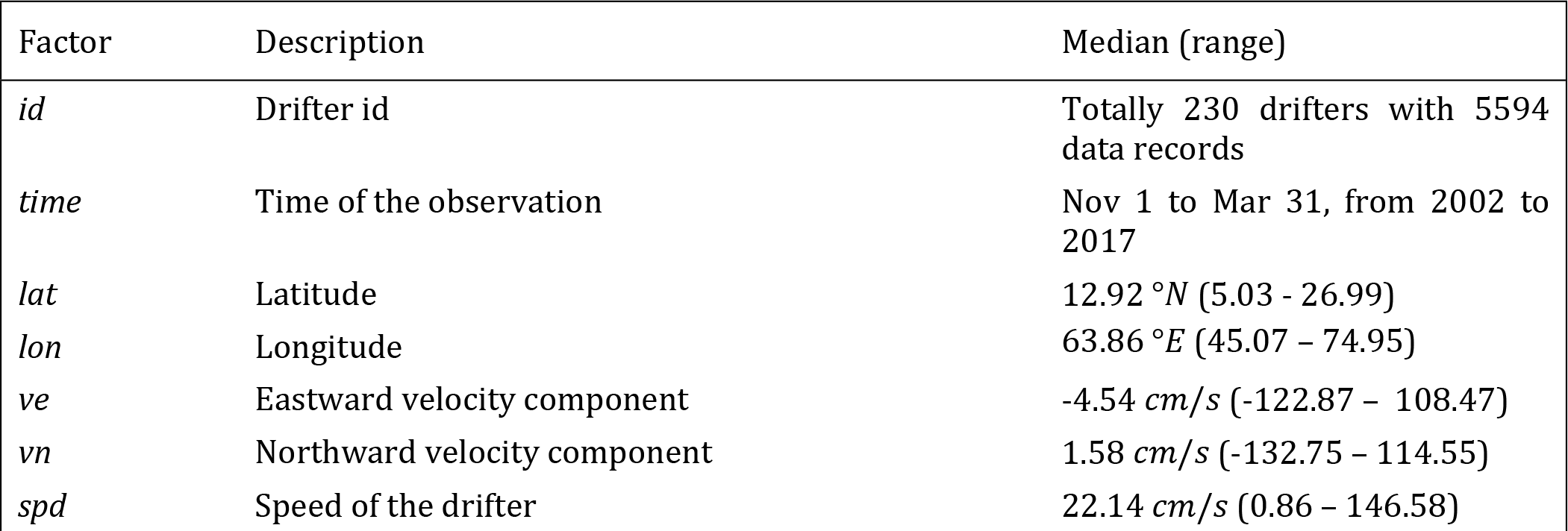
Factors used in the vLDS model.

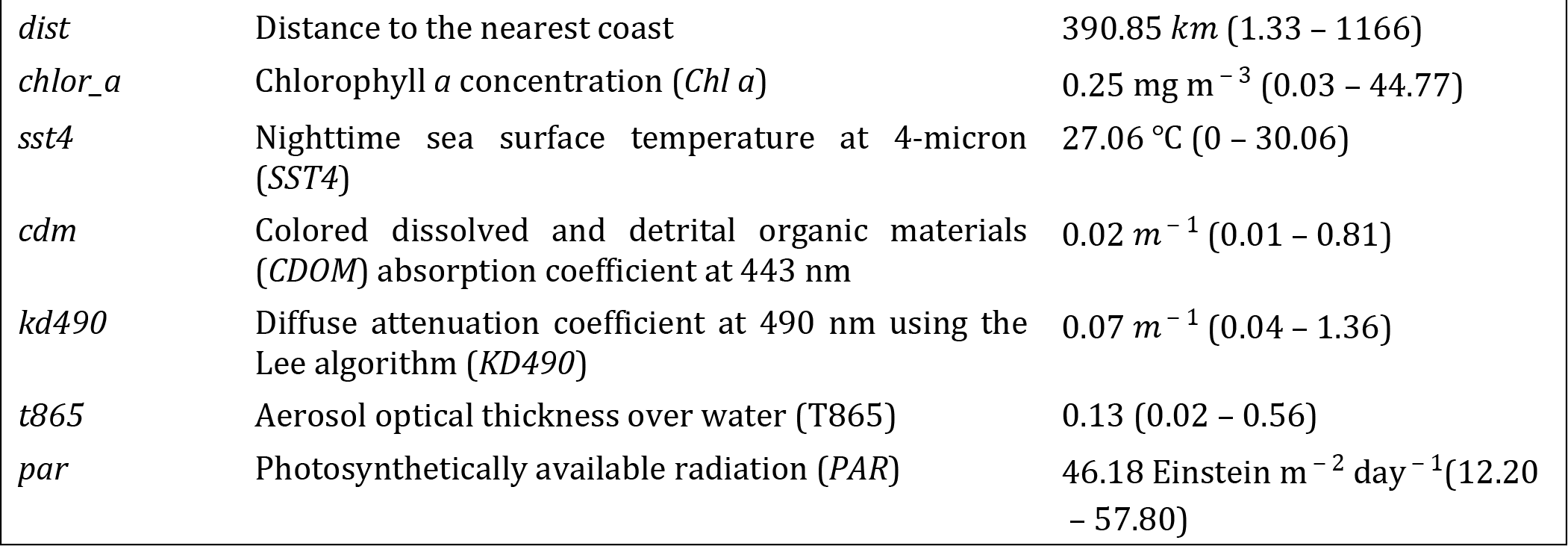

We note that each float record has information on its coordinates {*lat, lon, time*}.For convenience, we denote {*lat, lon, time*} as {*x, y, z*}, which is almost always not precisely on the grid of the Ocean Color dataset. To resolve this issue, we now describe the multidimensional interpolation operator to map the chlorophyll *a* concentration onto the GDP float dataset. For any float data point with coordinates {*x*_0_, *y*_0_, *t*_0_}, we identify the coordinate cube or the grid cell in the Ocean Color dataset that contains this point. Particularly, this cube has 8 vertices with coordinates generated by the outer product {*x*_*nearest*_, *x*_*next*_}⨀{*y*_*nearest*_, *y*_*next*_} ⨀{*t*_*nearest*_, *t*_*next*_}, where the subscripts *nearest* and *next* indicate the nearest and the furthest neighbours in the cube for each coordinate. We further denote the interpolation weight *w*_*x*_ for *x*_0_ as

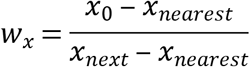

Similarly, we define the weights *w*_*y*_ and *w*_*t*_ for *y* and *t*. Using the function *f*(*x,y,t*) to represent the chlorophyll *a* concentration at the coordinate {*x,y,t*} we write the interpolated chlorophyll *a* concentration at {*x*_0_, *y*_0_, *t*_0_} as

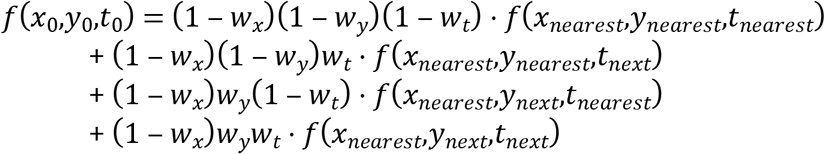

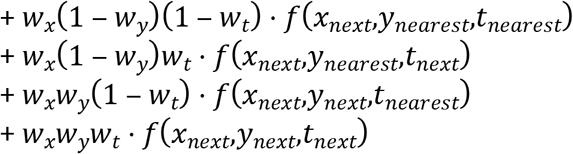

In our implementation, we have applied the interpolation process to the chlorophyll *a* concentration, distance to the nearest coast, and all other predictors. As an illustration of the interpolation process, we display the distance to the nearest coast [42] with a resolution of 4*km* in Fig 4A and the interpolated distance to the nearest coast for all the floats in Fig 4B.

**Fig 4.**
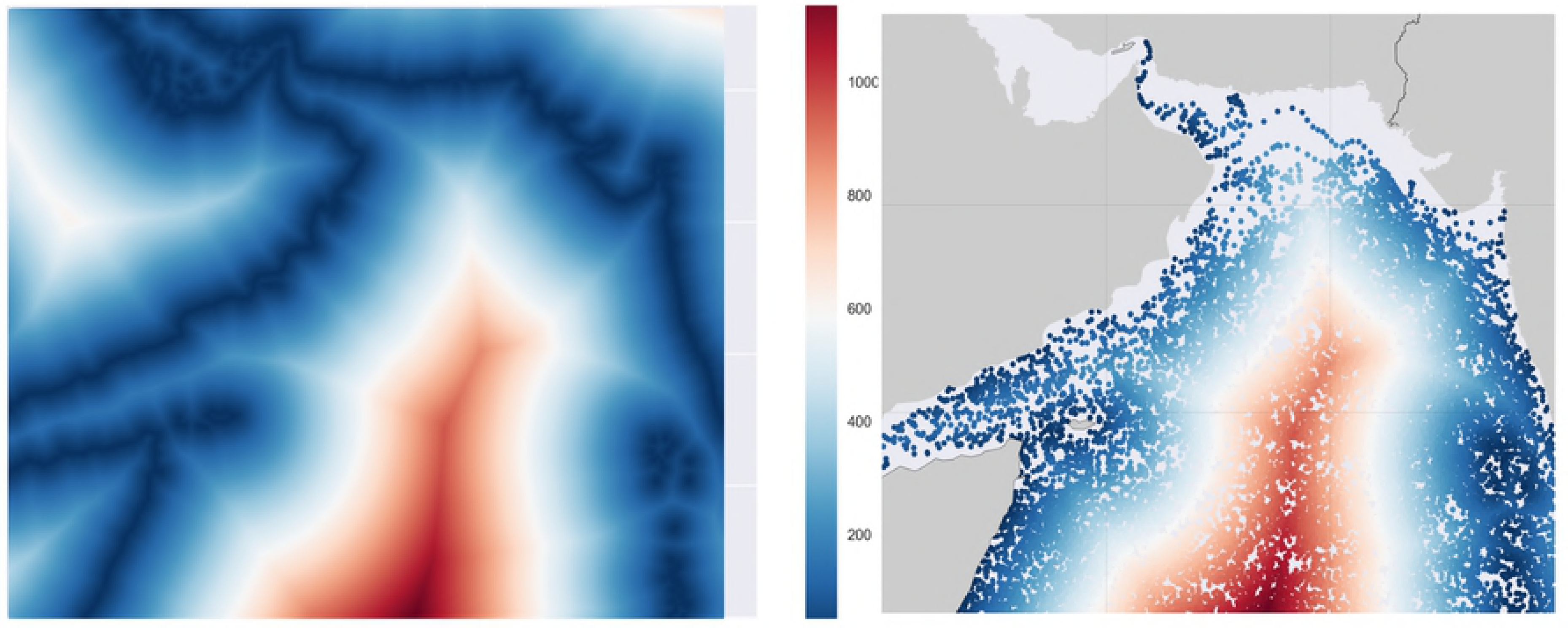
**(A)** Distance to the nearest coast for all the geographical locations in the Arabian Sea Region. **(B)** Interpolated values of the distance to the nearest coast for all the data points in the drifter dataset.

Using the multidimensional interpolation procedure described above, we map the satellite observations for each of the variables {*‘chlor_a’, ‘dist’, ‘cdm’, ‘kd490’, ‘t865’, ‘par’, ‘sst4’*} onto the GDP floats. Along with the information on the floats, namely {*‘time’, ‘id’, ‘lat’, ‘lon’, ‘ve’, ‘vn’, ‘spd’*}, the interpolated float dataset becomes high dimensional (Table 1). We remark that *‘chlor_a’* denotes the chlorophyll *a* concentration, *‘dist’* the distance from nearest coast, *‘cdm’* the colored dissolved and detrital organic materials (*CDOM*) absorption coefficient at 443 nm, *‘kd490’* the diffuse attenuation coefficient at 490 nm using the Lee’s algorithm (*KD490*), *‘t865’* the aerosol optical thickness over water (*T865*), *‘par’* the photosynthetically available radiation (*PAR*), *‘sst4’* the 4-micron nighttime sea surface temperature (*SST4*), *‘id’* the id of a float, *‘ve’* the eastward velocity component, *‘vn’* the northward velocity component, and *‘spd’* the speed of a float. These biological and physical factors are chosen to represent all the possible causes for the distribution of the *Noctiluca* blooms in the Arabian Sea. The *‘chlor_a’* measured during the period from November 1 to March 31 are mostly attributed to *Noctiluca* blooms. The *‘cdm’* measures *CDOM* the amount of dissolved organic materials in the sea water, which supports the growth of the *Noctiluca* [26]. The *‘dist’* measures the distance from the particle to the nearest coast, which is the source of the nutrient rich water. Moreover, during the winter season, the northwestern Arabian Sea off the coast of Oman experiences winter convective mixing (Nov-Jan), during which time nutrient rich, low-oxygen, cold water is brought to the surface both by convective mixing and by cyclonic eddy activity and benefit the growth of the *Noctiluca* in a complex and nonlinear fashion as described in the “Discussion & Conclusion” Section. The factor *‘par’* measures *PAR* the amount of light that is available on the sea surface for the photosynthesis of the symbiotic green algae (Fig 1B) living within *Noctiluca*. The *‘kd490’* provides an indication of the transparency of the water column and amount of light that penetrates into the sea water. The *‘t865’* represents the amount of particles in the atmosphere over the water, and is an indicator for the atmospheric deposition. The rest of the factors {*‘time’, ‘id’, ‘lat’, ‘lon’, ‘ve’, ‘vn’, ‘spd’*} are the spatio-temporal information describing the physical transport and dispersal of the *Noctiluca* blooms.

### Data preprocessing for vLDS

Due to the limitation of the satellite coverage mentioned in the “Data Collection” Section, there are missing values in each of the variables in the interpolated dataset. Our focus here is on the chlorophyll *a* concentration *‘chlor_a’*, since it is the key variable for *Noctiluca* blooms. The data structure of the post-processed float dataset is determined by the pre-processing steps for the variable *‘chlor_a’*.

The overall goal of data preprocessing is to keep the microscopic trajectory-based data structure intact, split the drifter trajectories that are spanning over multiple years, and remove the drifter trajectories that are too short for the learning algorithm, with the logical consideration of the physical and biological meaning. We consider that each period from November 1 to March 31 (during winter monsoon) represents one growth cycle of the *Noctiluca* bloom for a particular float or drifter. For any float with a unique id that has chlorophyll *a* data over two or more cycles, we split the data and assign a new derived float id to the data within each cycle, by adding a small increment 0.05 to the original float id. The resulting dataset allows the vLDS model to learn the microscopic dynamics of each cycle independently.

For each float id, we calculate the percentage of *‘NaN’* values in the dataset in the column *‘chlor_a’* and choose a threshold of 40%. There are two cases. First, if this percentage is smaller than the threshold, the data quality for this particular float is considered to be good, and we interpolate all the missing values for each of the variables in {*‘chlor_a’, ‘dist’, ‘cdm’, ‘kd490’, ‘t865’, ‘par’, ‘sst4’*}. See Fig 5 for an example, in which the time series is interpolated on *‘chlor_a’*. Also, after the interpolation process, there might still be gaps in the time series. For instance, the float might just do not have any record, including *‘NaN’*, in a certain short period. In this case, the float will be further split into continuous subseries. Therefore, every interpolated float time series will go through the second step, which we now describe, for checking and splitting.

**Fig 5.**
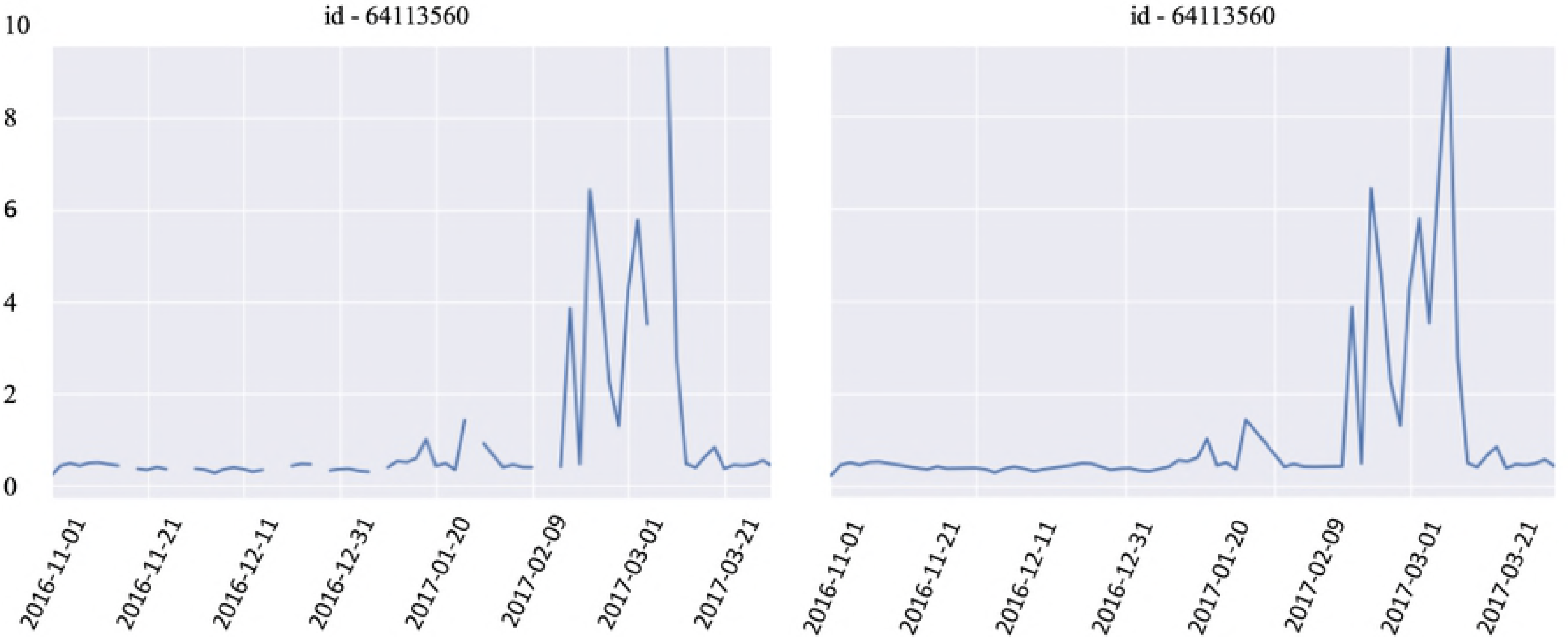
The time series of *‘chlor_a’* from the float id 64113560 on the left panel is interpolated and shown on the right panel.

In the other case, if this percentage of *‘NaN’* values is larger than the threshold, the data quality for this particular float is considered to unsuitable for interpolation. We split the discontinuous series into smaller continuous series on *‘chlor_a’*. We loop through the time series on *‘chlor_a’* and split it into smaller continuous series on *‘chlor_a’*. Moreover, we assign a new derived float id to each newly generated shorter series, by adding a small increment 0.03 to the original float id. Also, we drop any series on *‘chlor_a’* of length 1. See Fig 6 for an example, in which we split the time series on *‘chlor_a’* into 5 different shorter series, and we drop three series of length 1.

**Fig 6.**
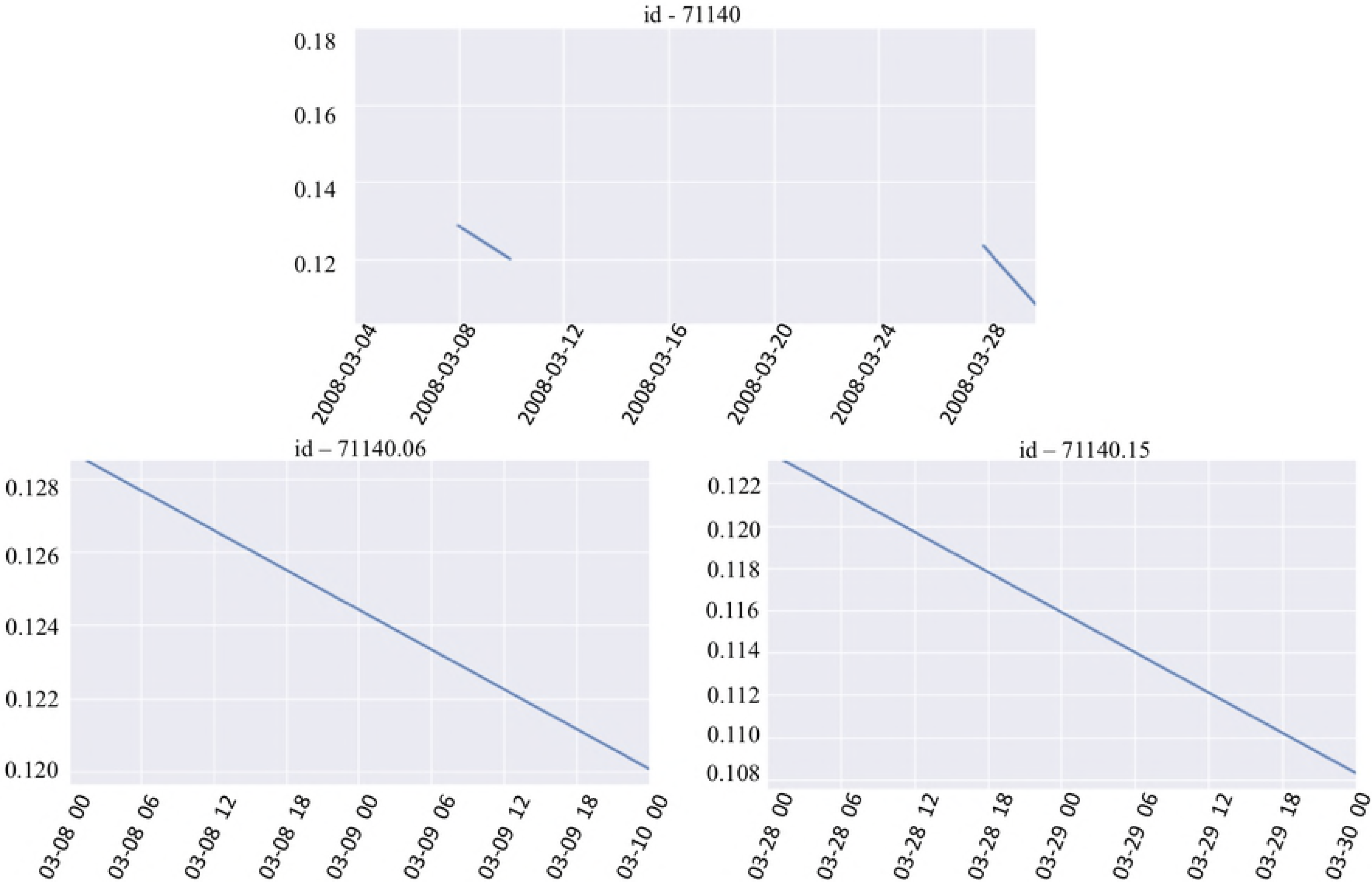
The time series of *‘chlor_a’* from the float 71140 on the top panel is split into 5 different shorter series. Three series of length 1 are dropped, and the remaining two are shown on the bottom panel

### Linear Dynamical Systems (LDS)

After being preprocessed, the merged GDP floats dataset consisted of 186 float records. For each float, the measurement is a multivariate time series ***y***, given by

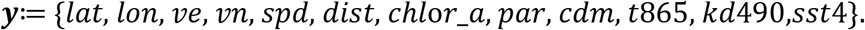

We use 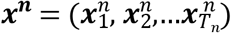 to denote the latent variables and 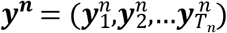 the observations from the float *n* and *T*_*n*_ for the length of the time series of the float *n*. The plain version Linear Dynamical System (LDS) is an adaptive procedure that can learn from data to recover the latent relationship between ***y*** and ***x***, using assumptions of linear relationships at time *i* between ***x***_***i***_ and ***x***_***i*-1**_, ***y***_***i***_ and ***x***_***i***_, with Gaussian Noise. For the moment, we omit the superscript *n* and focus on one float. More specifically, we assume that for a specific float the time series of the latent variables and the observations has the following relationships

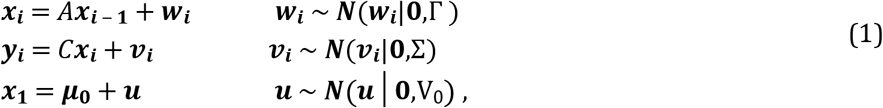

where ***w***_***i***_, ***v***_***i***_, ***u*** are noise terms. The LDS model fits the model parameters Θ: = {*A,C*,Γ,Σ, **μ**_0_, V_0_} by taking the expectation over the latent variables ***x***_***i***_|*θ*_***old***_, and maximizing the log-likelihood of the complete data {***x,y***|***θ***}, where ***θ***_***old***_ is the model parameter from the previous iteration and ***θ*** is the parameter that we are seeking at the current iteration. In the expectation step, with the parameter ***θ***_***old***_, the mean and the variance of the posterior marginal latent variables ***x***_***i***_|***θ***_***old***_, ***y***_1_, ***y***_2_,…***y***_***i***_ at time ***i*** (see float 1 in Fig 7A) and the mean and the variance of the posterior marginal latent variable ***x***_***i***_| ***θ***_***old***_, ***y***_**1**_, ***y***_**2**_,…***y***_***T***_ based on the information at all time (see float 1 in Fig 7B), are calculated using the forward and backward iterations. Here, *T* is the total length of a particular time series on a float. These marginal variables lead to the sufficient statistics of the complete data{***x***, ***y***,|***θ***, namely the 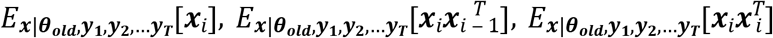. Using these sufficient statistics [54, 55], we obtain the updated LDS model parameters **Θ**: = {*A,C*,Γ,Σ, **μ**_**0**_, V_0_}. The graphical model [56] of the plain LDS is schematically plotted in Figs. 7A, 7B as one branch, for instance, the branch of float 1. The workflow of the plain LDS model is described in Algorithm 1.

**Fig 7.**
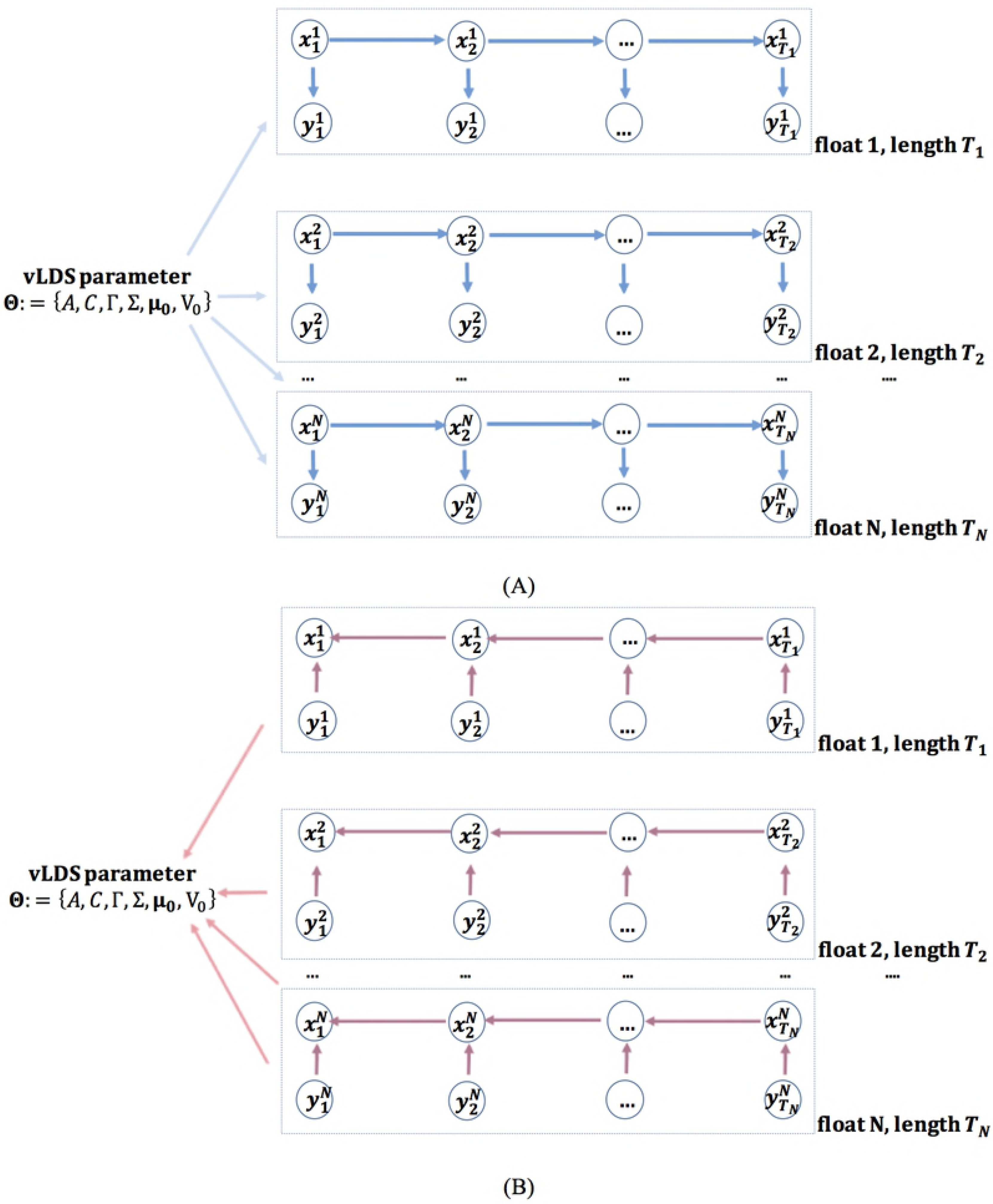
**(A)** Information flow of the forward iteration in the Expectation step of the vLDS model for computing the mean and the variance of the posterior marginal latent variables ***x***_***i***_|***θ***_***old***_, ***y***_**1**_, ***y***_**2**_,…***y***_***i***_ at time ***i***. **(B)** Information flow of the backward iteration in the Expectation step of the vLDS model for computing the mean and the variance of the posterior marginal latent variables ***x***_***i***_|***θ***_***old***_, ***y***_**1**_, ***y***_**2**_,…***y***_***T***_ based on information at all time 1…*T*.

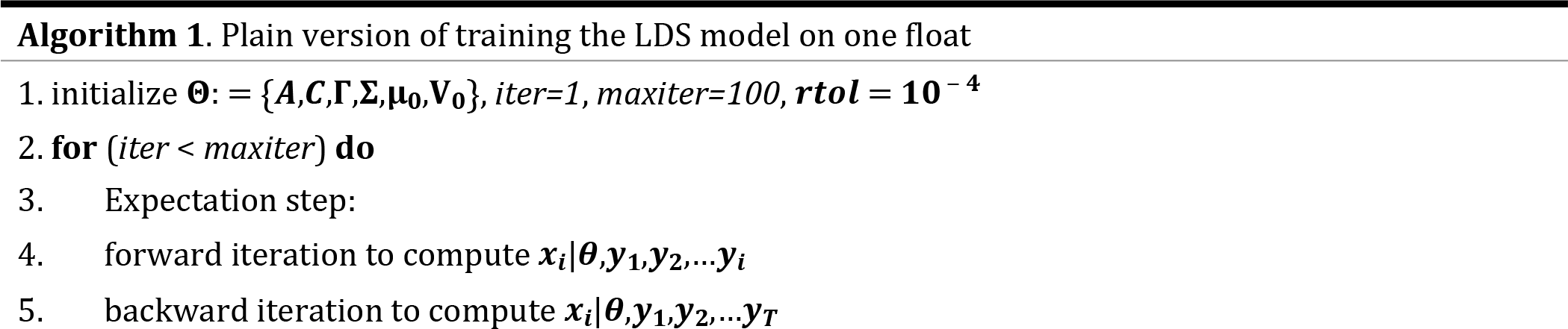

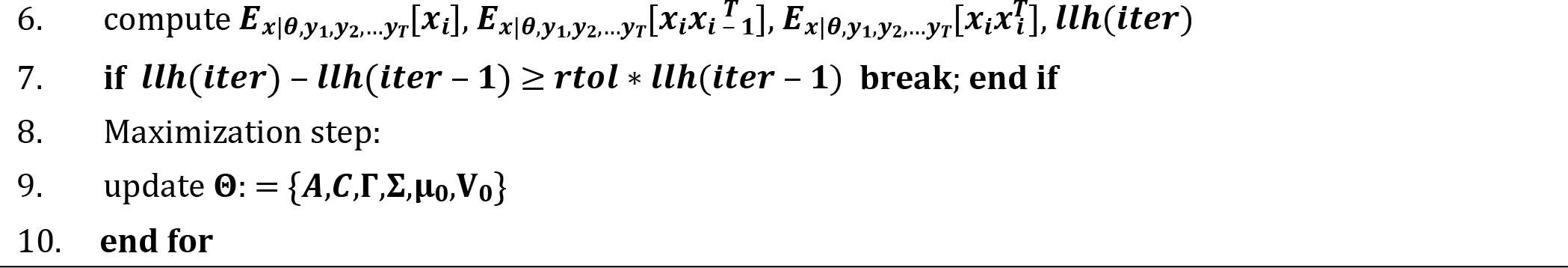

### Variable-length Linear Dynamical Systems (vLDS)

In this study, each float generates one or more statistically independent time series of the *Chl a* concentration, due to the interpolation or splitting process discussed in the “Data Preprocessing” Section. For the preprocessed dataset with 186 floats, we treat it as multiple multivariate time series, each with a unique id. Also, we note that the lengths of the time series in the dataset are mostly different, due to the irregularity in the longevity of the floats. The variable-length Linear Dynamical Systems model is specifically designed for this situation, and it summarizes and recovers the latent dynamics from multiple multivariable time series with a different time span.

To fit the vLDS model, we start with some initial parameter **Θ**_**0**_, which is shared across all floats in the dataset. We keep the superscript *n* here. The Expectation step is carried out on each float id, using a two-loop forward and backward smoothing step to compute the conditional expectations of the sufficient statistics of the complete data {***x**, **y***|***θ***}, namely the 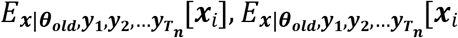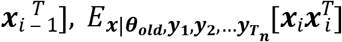. In the maximization step, we use the averaging formula derived in Equation (5) across all floats to update the model parameter **Θ**: = {*A,C*,Γ,Σ, **μ**_**0**_, V_0_}.

We emphasize that for a particular drifter *n*, the multivariate observation 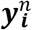, *i* = 1,2,3,…*T*_*n*_, contains all the physical and biological information along a drifter trajectory at time *i* and it is the multivariate latent random variable 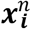 that we are solving for from the vLDS model to represent the hidden dynamics between different components of 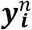, namely,

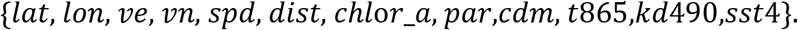

It is possible that some of the components of 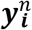, for instance, the *t*865 in our study, as demonstrated in the “Discussion & Conclusion” Section, are not much involved in the latent dynamics. Therefore, the dimension of the latent space recovered by the latent variable 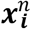 might be smaller than the dimension of the observations 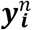. In this study, the latent dimension in 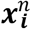, as determined by the cross-validation procedure, turns out to be 11, and the dimension of the observations 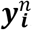 is 12.

To test our benchmarked hypotheses in the “Background” Section, we first generate the predicted values of 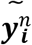 by using the equation (1) with the recovered latent variables 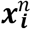 from the vLDS model.

Since the vLDS model automatically maximizing the log-likelihood of the complete data, including the latent variable ***x*** and the observational variable ***y***, it is beneficial to visualize the predictive plots of all predictors along the drifter trajectories, as shown in the “Discussion & Conclusion” Section, and study the performance metric R-squared, as described in the “Results” Section.

### Probabilistic computation of vLDS

We assume that the observed multivariate variable 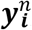, including the chlorophyll *a* concentration, of each float is evolving independently with other floats, except that they are driven by the same underlying biological and physical forces. It is this assumption that allows the vLDS model to share the same model parameters **Θ**: = {*A,C*,Γ,Σ, **μ**_**0**_, V_0_} in all branches in Fig 7A, 7B and makes the vLDS model a powerful algorithm to summarize and capture the population-level structures along drifter trajectories. Multiple variants of the LDS model have been applied successfully at the microscopic scale in several research areas such as computational neuroscience [19–20, 57–58] and sound tracking [22].

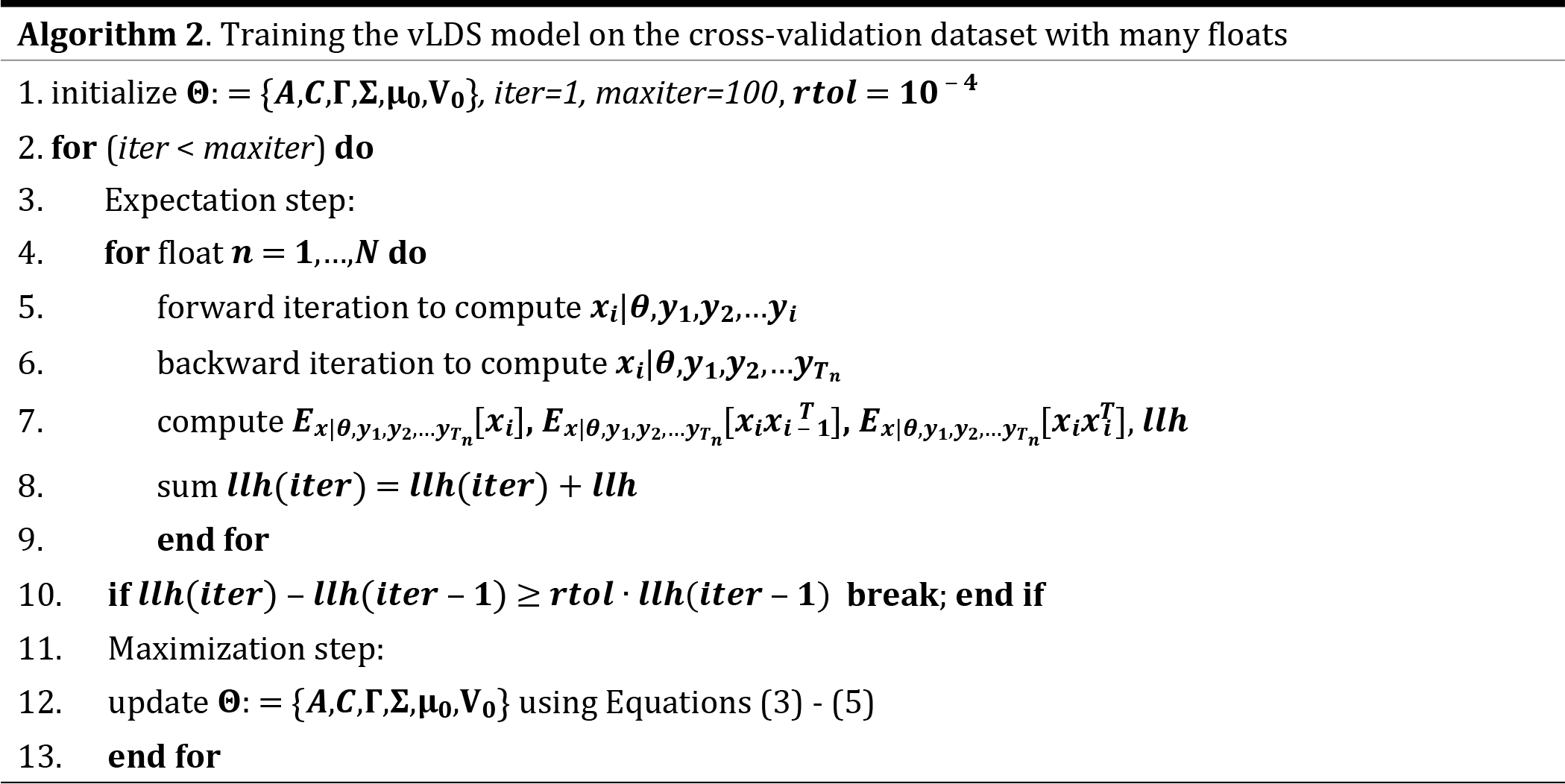

Although the distribution of the complete data {***x**, **y***|***θ***} depends on the model parameter ***θ***, we omit the dependence on ***θ*** for notational ease in our following derivation. For a particular float *n*, letting *i* be the time step and *T*_*n*_ be the total length of the time series, the distribution of the complete data (namely the observations ***y*** and the latent variables ***x***), can be written as

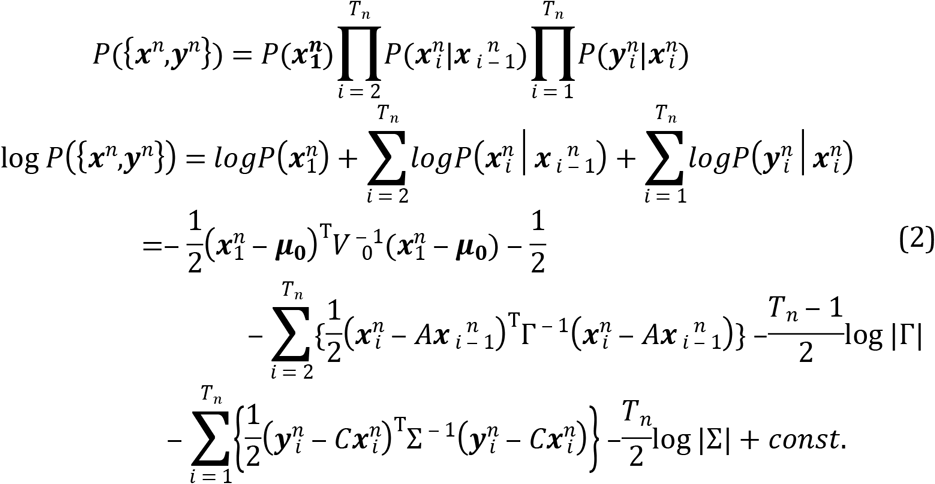

By the assumption of independence, the joint probability of the observations and state variables across all floats expands into the product of the joint probability of the observations and state variables of all the time series generated by each float *n* = 1,2…*N*

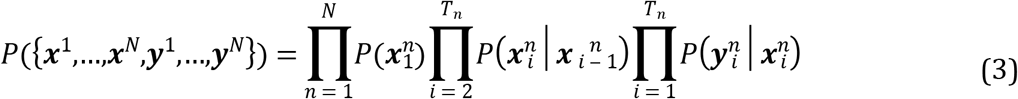

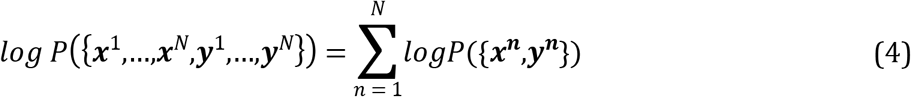

We note that, for each float, the preprocessed data of this float might generate multiple multivariate time series (see the “Data Preprocessing” Section for more details.) Under the traditional i.i.d. assumptions, the objective function of the vLDS model is simply the addition of the objective functions for each individual time series' log-likelihood with all the parameters **Θ**: = {*A,C*,Γ,Σ, **μ**_**0**_, V0} that define the plain version LDS for each float (or each box-branch in Fig 7) being shared across all the floats. Using the Expectation-Maximizing algorithm, the Expectation step can be carried out using a two loop backward and forward iteration for each time series independently, due to the conditional independence of the state variables across different time series of different floats. However, in the maximization step we need to average the vLDS model parameter across all the time series from all the floats.

The derivation of the update formula for the vLDS model parameters **Θ**: = {*A,C*,Γ,Σ, **μ**_**0**_, V_0_} follows directly by taking derivatives of the complete data log-likelihood with respect to each component in **Θ** and by using the standard results from the Maximum Likelihood Estimators of the mean and variance for the Gaussian Distribution. For a dataset of many floats, the complete data log-likelihood has an addition summation sign running through *n* = 1, 2…*N* in Equation (4). We use *T*_*n*_, instead of *T*, to denote the length of the time series of the float *n*. Given the fact that the derivative of a linear combination of functions is a linear combination of derivatives of each function, we can write the updating formula for the maximization step as:

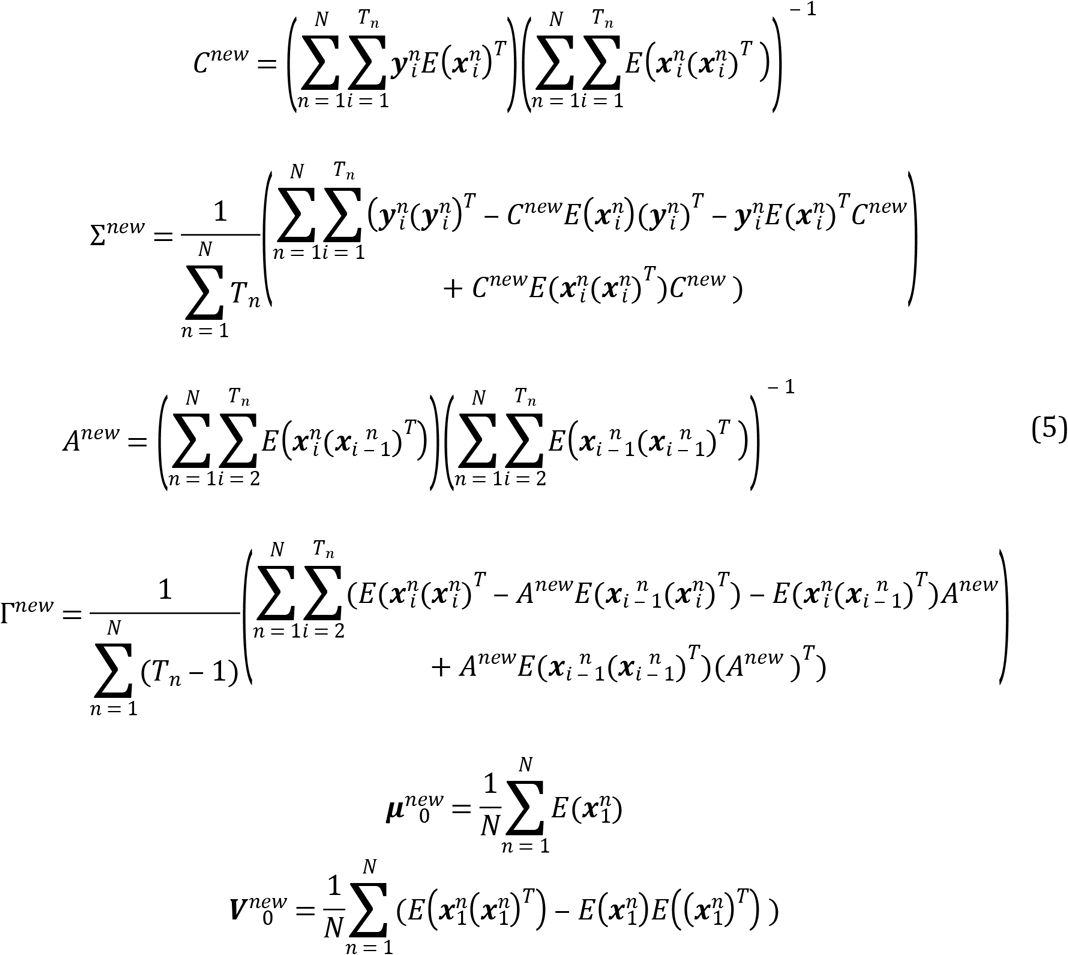

The workflow of the vLDS model is described in Algorithm 2, and the averaging of the vLDS model parameters **Θ** across all the floats in equation (4) is carried out in step 12. During this procedure, the vLDS model adaptively learns the latent dynamics of the underlying process. We have fitted the chlorophyll *a* concentration for the particular float in Fig 5 with id 64113560. The result is displayed in the “Discussion and Conclusion” Section.

Each Expectation-Maximization cycle of the LDS model for Gaussian random variables is guaranteed to increase the value of the complete data log-likelihood. Therefore, a standard stopping criteria for the Expectation-Maximization algorithm is based on the complete data log-likelihood in Equations (2), (4) with a relative tolerance *rtol* = 10 ^−4^ and maximum iteration 100.

One of the key model parameters in the LDS modeling is the dimension *k* of the latent space, namely, the number of components in the latent variable ***x***. It is the dimension of the subspace generated by the projection of the full feature space onto the latent subspace, whose projection back onto the full feature space in Equation (1) under the vLDS linear transformation matrix *C* maximizes the complete data log-likelihood. A larger *k* indicates that there are more independent factors in the latent space of ***x*** driving the underlying dynamical system of {***x**, **y***}. Moreover, varying the values of the dimensionality *k* induces a family of different vLDS models (1) - (3) indexed by *k*. To select the model with the most appropriate parameter *k*, we carry out a 10-fold cross-validation [59–61] on the parameter *k* and choose the optimal *k* that achieves the maximum complete data log-likehood on the test dataset. More specifically, we group the dataset by float ids. We hold a portion of the floats ids and consider them as the heldout dataset. We take the rest of the float ids as the cross-validation dataset. In the cross-validation step, we split the cross-validation set evenly into 10 folds. Each time we take one fold as the testing dataset, we take the rest as the training dataset. We fit the vLDS parameter **Θ**: = {*A,C*,Γ,Σ, **μ**_**0**_, V_0_} on the training dataset and compute the complete data log-likelihood on the testing dataset using this newly fitted parameter **Θ**. The complete data log-likelihood is averaged for different testing fold for a fixed *k*. Then, we repeat the process for different values of *k*. See Table 2 and Fig 8 for the complete data log-likelihood generated by different cross-validation trails. The averaged complete-data log-likelihood across different testing set is maximized by *k* = 11.

**Table 2.** 10-fold cross-validation on the parameter *k*. The optimal *k* that achieves the maximum complete data log-likehood on the test set is *k* = 11. The unit of the test-dataset log-likelihood in the table is 10^4^.

|  | k | 1 | 2 | 3 | 4 | 5 | 6 | 7 | 8 | 9 | 10 | 11 | 12 |
| --- | --- | --- | --- | --- | --- | --- | --- | --- | --- | --- | --- | --- | --- |
| fold | $ll_{test}$ | | | | | | | | | | | | |
| 1 |  | -1.44 | -1.36 | -1.33 | -1.23 | -1.15 | -1.19 | -1.15 | -1.04 | -1.14 | -1.18 | -1.01 | -1.65 |
| 2 |  | -1.33 | -1.22 | -1.20 | -1.09 | -0.98 | -1.02 | -0.95 | -0.83 | -0.94 | -0.97 | -0.81 | -1.42 |
| 3 |  | -1.58 | -1.53 | -1.47 | -1.27 | -1.25 | -1.19 | -1.26 | -1.10 | -1.19 | -1.15 | -1.02 | -1.46 |
| 4 |  | -1.29 | -1.18 | -1.15 | -1.04 | -0.98 | -0.97 | -0.94 | -0.82 | -0.88 | -0.86 | -0.80 | -1.11 |
| 5 |  | -1.44 | -1.33 | -1.30 | -1.17 | -1.12 | -1.11 | -1.10 | -0.91 | -1.06 | -1.11 | -0.88 | -1.57 |
| 6 |  | -1.58 | -1.45 | -1.45 | -1.34 | -1.22 | -1.36 | -1.31 | -1.17 | -1.27 | -1.19 | -1.14 | -1.70 |
| 7 |  | -1.71 | -1.55 | -1.52 | -1.34 | -1.25 | -1.22 | -1.19 | -1.02 | -1.07 | -1.09 | -0.95 | -1.31 |
| 8 |  | -1.14 | -1.02 | -1.02 | -0.93 | -0.84 | -0.88 | -0.87 | -0.73 | -0.76 | -0.76 | -0.71 | -0.95 |
| 9 |  | -1.63 | -1.48 | -1.44 | -1.27 | -1.17 | -1.18 | -1.19 | -0.99 | -1.06 | -1.10 | -0.94 | -1.35 |
| 10 |  | -1.52 | -1.40 | -1.35 | -1.22 | -1.20 | -1.13 | -1.11 | -0.92 | -0.99 | -1.04 | -0.85 | -1.33 |
| Average |  | -1.47 | -1.35 | -1.32 | -1.19 | -1.11 | -1.12 | -1.11 | -0.95 | -1.04 | -1.05 | **-0.92** | -1.38 |

**Fig 8.**
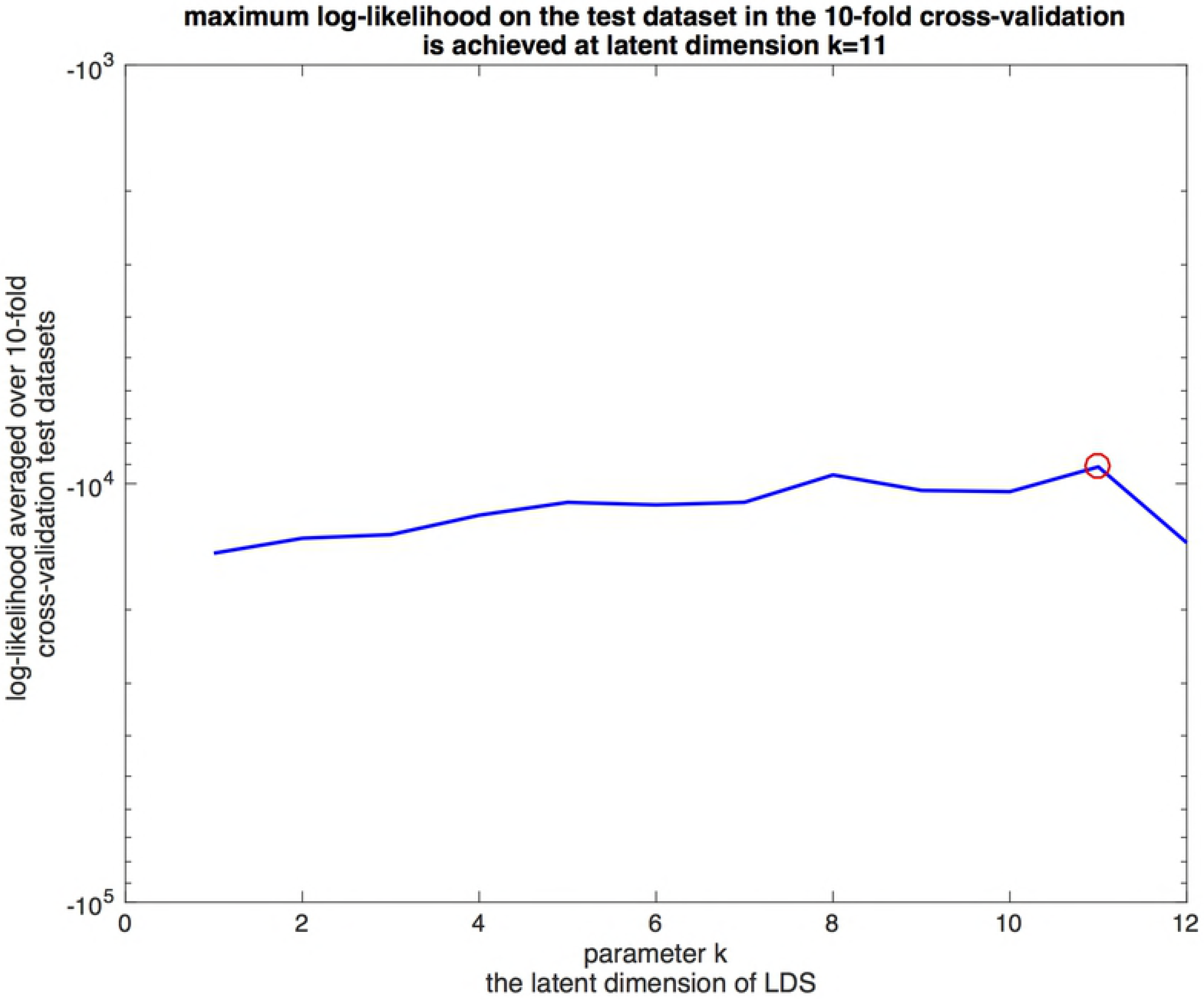
10-fold cross-validation on the parameter *k* and the optimal *k* that achieves the maximum complete data log-likehood on the test set is *k* = 11.

With the optimal value *k* = 11 of the latent space dimension identified, we fit the vLDS model one more time with the full cross-validation dataset to generate the vLDS model parameter. In Fig 9, we display the log-likelihood convergence of the Expectation-Maximization algorithm for the complete cross-validation dataset and five individual floats in the cross-validation dataset. We note that from Equations (2) - (4), the log-likelihood of the complete cross-validation dataset is the sum of the log-likelihood of each individual floats in the cross-validation dataset (step 8 in Algorithm 2).

**Fig 9.**
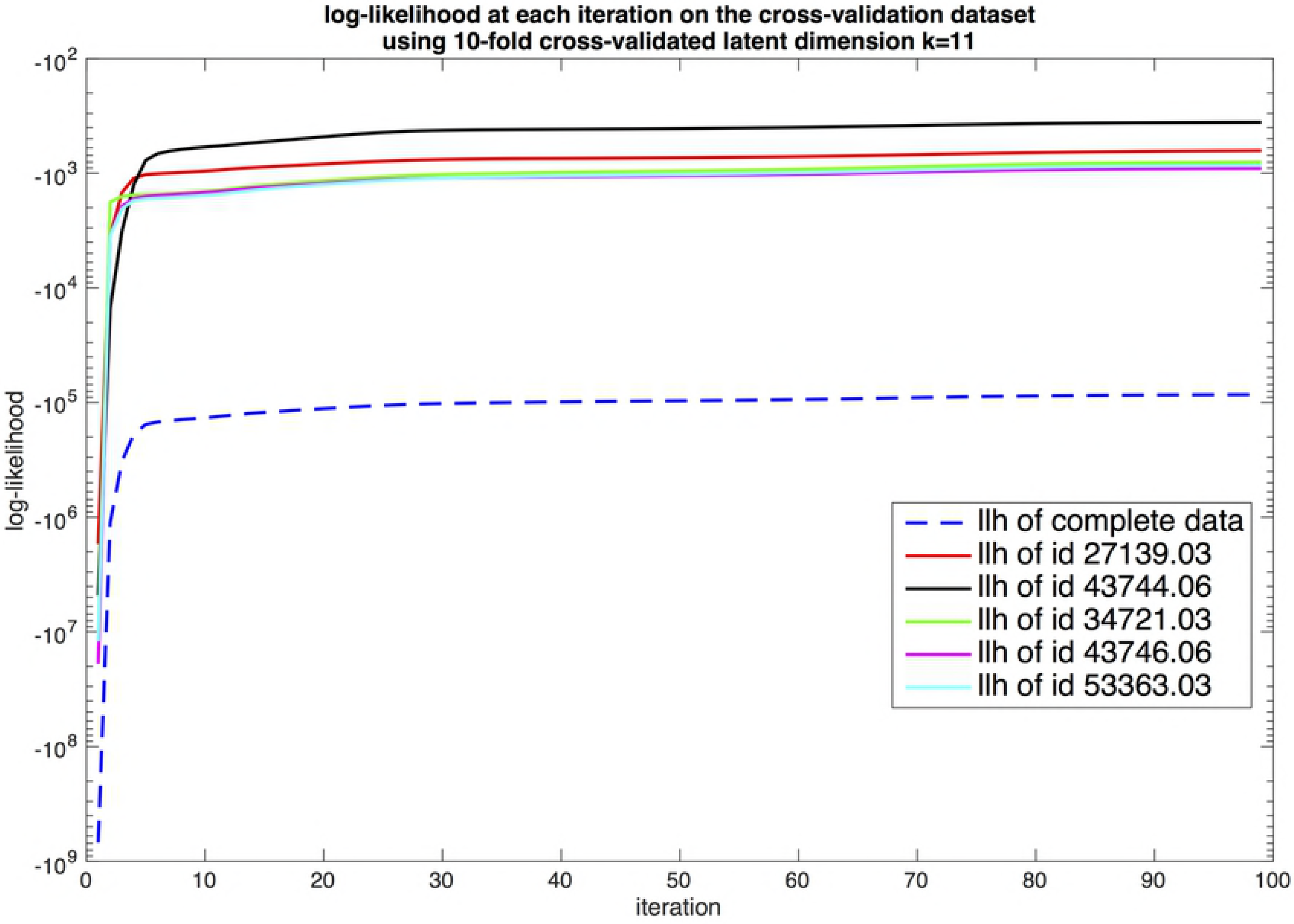
Convergence history of the log-likelihood of the complete cross-validation dataset and a sample of the convergence history for 5 floats.

## Results

With the optimally selected latent space dimension = 11, the vLDS algorithm obtains a set of model parameters **Θ**: = {*A,C*,Γ,Σ, **μ**_**0**_, V_0_} when the stopping criteria inside the Expectation-Maximization algorithm is reached. In Fig 10, we display the prediction results for some drifter ids in the cross-validation dataset, using Equation (1) and the expected conditional mean of the latent variables at the last iteration of the Expectation steps 4, 5, and 6 in Algorithm 2. The dark lines are the observations, and the cyan lines are the predictions. Most of the hidden dynamics of the float profiles inside the cross-validation dataset are well captured by the vLDS model. We note the positive correlations among *‘chlor_a’, ‘cdm’*, and *‘kd490’* in the recovered vLDS latent dynamics (cyan lines in Fig 10) at the microscopic population-level for all drifters in the cross-validation dataset. The model captures this correlation with some overshooting or undershooting in certain regions. Also, *‘t865’*, the aerosol optical thickness over water, turns out to be independent of the chlorophyll *a* concentration and other ocean profiles. Moreover, the spatial information, namely, the longitude, latitude, velocity, speed of the float, and distance from the nearest coast, is all well recovered by the vLDS model (*lat* and *spd* are not shown in Figs 10, 11 due to space limitations).

**Fig 10.**
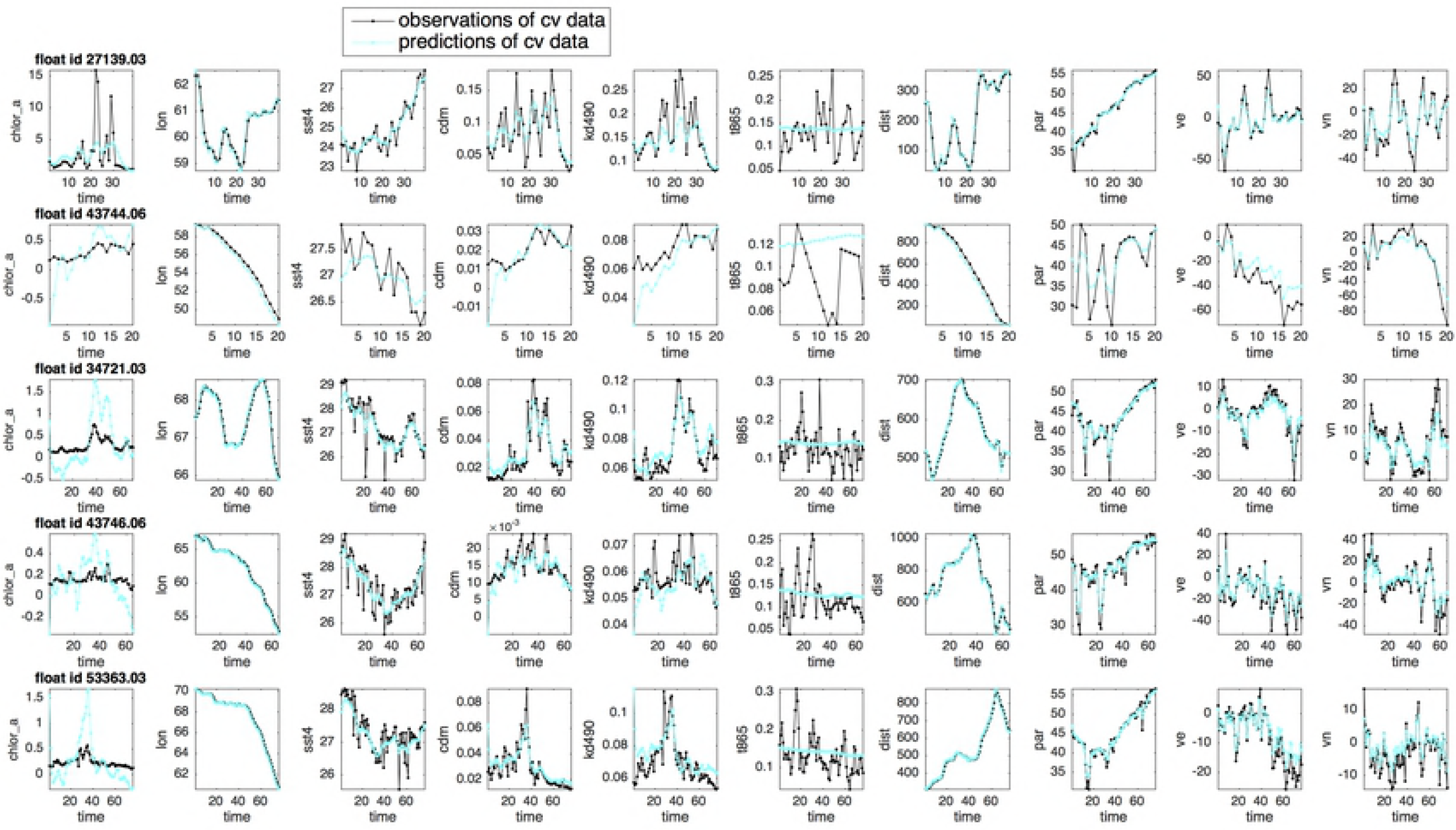
Predictions of the drifter profiles for the floats in the cross-validation dataset, using the expected conditional mean of the latent variables at the last iteration of the Expectation-Maximization algorithm.

**Fig 11.**
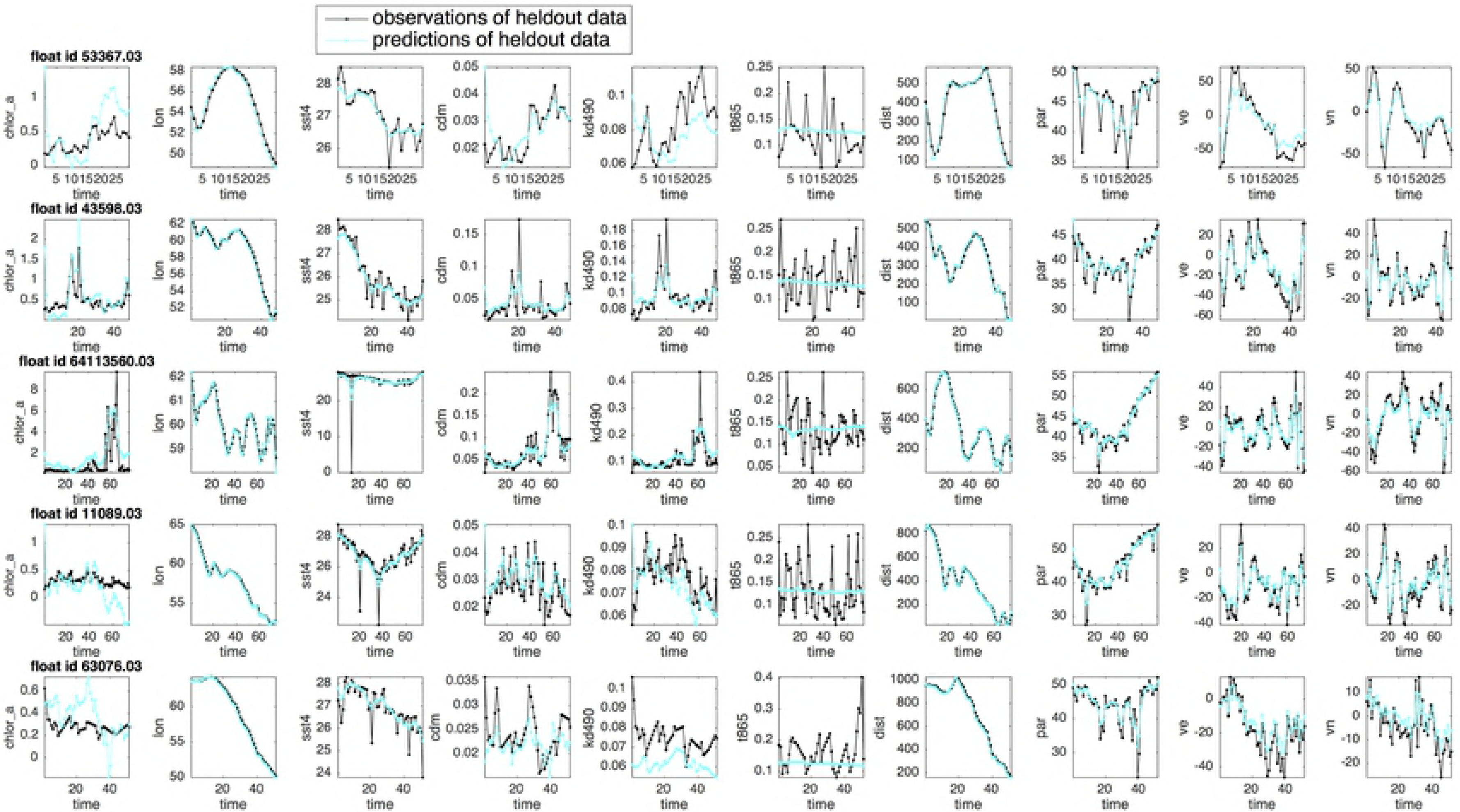
Predictions of the drifter profiles for the floats in the heldout dataset, using the expected conditional mean of the latent variables generated by one iteration of the forward-backward smoothing process of the Expectation-Maximization algorithm with the vLDS model parameter **Θ**: = {*A,C*,Γ,Σ, **μ**_**0**_, V_0_} optimized on the cross-validated dataset.

We next examine on the robustness of the vLDS model. The floats in the heldout dataset are not used in the cross-validation process or the model’s parameter estimation process. Therefore, the heldout dataset is totally unknown to the vLDS learning algorithm. We use the cross-validated latent dimension *k* = 11, and the model parameter **Θ**: = {*A,C*,Γ,Σ, **μ**_**0**_, V_0_} generated by training the vLDS model on the cross-validation dataset. Applying one iteration of the forward-backward smoothing process, namely, merely one iteration of the Expectation steps 4, 5, and 6 in Algorithm 2, to each float in the heldout dataset, we obtain the predictions of their profiles (Fig 11). Most of the hidden dynamics along drifter trajectories for the floats in the heldout dataset, which is totally unknown to the learning algorithm, is well captured by the vLDS model. It clearly demonstrates the generalizability of the vLDS model’s capability to summarize and capture the population-level structures along the drifter trajectories on unknown datasets.

We note again the positive correlations among *‘chlor_a, ‘cdm’*, and *‘kd490’* in the recovered vLDS latent dynamics (cyan lines in Fig 11) at the microscopic population-level for the drifters in the heldout dataset. Even in this heldout dataset, the model is capable of capturing this correlation with some overshooting or undershooting in certain regions. Again, *‘t865’*, the aerosol optical thickness over water, seems to be independent of the *Chl a* concentration and other ocean profiles in the heldout dataset. The vLDS model simply estimate the mean value for the variable ‘t865’ in both the cross-validation and heldout datasets. However, by comparing the predictions generated by the vLDS model for ‘t865’ and for other variables, we conclude that the variable *‘t865’* is not much involved in the latent dynamics of the *Noctiluca*’s growth. Otherwise, the predicted values of *‘t865’* should match its observations in Fig 10, 11, and the *R*^2^ of *‘t865’* in Table 3 should not be too small. Evidently, the vLDS model provides us the opportunity to verify the causal relationships of all variables at the microscopic population-level along the drifter trajectories, a feature the must be lacking in any models that do not take the trajectory-based population-level structure into consideration. Moreover, the spatial information of the heldout floats, namely, longitude, latitude, velocity, speed of the float, and distance from the nearest coast, is all well recovered by the vLDS model.

To quantify the performance of the vLDS model, we use the R-squared metric (*R*^2^). In Table 3, the total sum of squares (SSTotal), sum of squared errors of predictions (SSE), and *R*^2^, which is the portion of the variance explained by the predictive model, are computed for both the cross-validation and heldout datasets. Although the vLDS model has the log-likelihood of the complete data in Equations (3), (4) as its own performance metric, we use *R*^2^ here for an intuitive interpretation. The quantitative results reflect the visualization in Fig 10, 11. The feature *‘t865’* has a very small value in its *R*^2^ metric and does not exhibit any predictive power. The vLDS recovers the spatio-temporal information well, and explains most of the variance in the biological factors {*‘cdm’, ‘kd490’, ‘par’, ‘sst4’, ‘chlor_a*}.

**Table 3.** R-squared (*R*^2^) metric for the cross-validation dataset and heldout dataset. The *R*^2^ is computed for each individual feature and for all the features aggregated together.

|  | cross-validation data |  |  | heldout data |  |  |
| --- | --- | --- | --- | --- | --- | --- |
| | SSTotal | SSE | $R^2$ | SSTotal | SSE | $R^2$ |
| *lat* | 1.17E+05 | 3.78E+02 | 0.99 | 4.32E+03 | 1.86E+01 | 0.99 |
| *lon* | 2.32E+05 | 4.42E+02 | 0.99 | 3.52E+03 | 3.23E+01 | 0.99 |
| *ve* | 2.37E+06 | 3.67E+05 | 0.85 | 1.51E+05 | 2.49E+04 | 0.84 |
| *vn* | 2.36E+06 | 4.51E+05 | 0.81 | 1.07E+05 | 1.63E+04 | 0.85 |
| *spd* | 1.78E+06 | 2.62E+05 | 0.85 | 7.20E+04 | 1.09E+04 | 0.85 |
| *dist* | 3.55E+08 | 2.05E+06 | 0.99 | 1.78E+07 | 1.52E+05 | 0.99 |
| *cdm* | 1.65E+01 | 1.88E+00 | 0.89 | 3.40E-01 | 4.00E-02 | 0.87 |
| *kd490* | 2.43E+01 | 8.02E+00 | 0.67 | 3.19E-01 | 1.39E-01 | 0.56 |
| *t865* | 1.34E+01 | 1.29E+01 | 0.04 | 7.30E-01 | 7.49E-01 | -0.04 |
| *par* | 2.41E+05 | 1.80E+04 | 0.93 | 9.43E+03 | 7.87E+02 | 0.91 |
| *sst4* | 1.20E+04 | 2.07E+03 | 0.83 | 1.02E+03 | 5.13E+02 | 0.49 |
| *chlor_a* | 4.60E+04 | 2.19E+04 | 0.52 | 2.59E+02 | 1.46E+02 | 0.43 |
| Aggregated | 3.62E+08 | 3.17E+06 | 0.99 | 1.82E+07 | 2.06E+05 | 0.98 |

## Discussion & conclusion

We have introduced a new model vLDS and showed it offers a new approach to recover microscopic biogeochemical mechanisms underlying chaotic drifter trajectories that might be unobservable at macroscopic scale or accessible only in controlled laboratory experiments. The vLDS model generates predictions that recover the causal relationship among the *Noctiluca* blooms, physical dispersal, and biological environments (Fig 10, 11, Table 3.) The model’s generalization capability also summarizes, recovers, and predicts the latent dynamics from unknown heldout datasets, thus inspiring confidence in our microscopic and macroscopic findings. The highly correlated relationships between the *‘chlor_a’* and *‘cdm’* (colored dissolved organic matter *CDOM*), and between the *‘chlor_a’* and *‘kd490’* (light under the sea surface) are close to linear. The tightly correlated relationships between the *‘chlor_a’* and *‘par’* (light on the sea surface *PAR*), and between the *‘chlor_a’* and *‘sst4’* (sea surface temperature *SST4*) are nonlinear. The atmospheric deposition *‘t865’* is not involved in the underlying dynamics of the *Noctiluca* blooms.

These results confirm our hypotheses in the “Background” Section from the microscopic perspective, and confirms the impact of both the physical transport and biological factors of light, and nutrients, proxies for the latter being *CDOM* [26, 62–65], on the distribution of the *Noctiluca* blooms. Also, from the test results, the nutrient and light are two important positive factors for the *Noctiluca*’s growth and that the atmospheric deposition does not contribute much to this process. More specifically, the nutrient-rich water flowing into sea are increasing productivity and expanding the hypoxia zone as oxygen gets depleted. Since the atmospheric deposition measured by *T865* does not contribute much to the source of nutrient (Fig 10, 11, Table 3), the potential sources of the organic matter are including the domestic and industrial outfall from countries bordering the Arabian Sea, and the Northeast Monsoon (NEM, Nov-Jan) and winter convective mixing of deep nutrient rich, oxygen poor waters. As plankton grows, it uses up oxygen in the water. Falling oxygen levels allow *Noctiluca* to outcompete diatoms during winter. It has also been confirmed from the microscopic dynamics recovered by the vLDS Model that *Noctiluca* grow faster in lighted than in dark areas on the sea surface and in the sea water.

We have demonstrated the effectiveness of the vLDS model as a microscopic statistical modeling approach for detecting important causal relationships in biogeochemical processes. Although the trajectories of the oceanographic probing devices are chaotic and the dataset is high dimensional, the vLDS model is very parsimonious on model parameters. The model only requires **Θ**: = {*A,C*,Γ,Σ, **μ**_**0**_, V_0_} and the latent-space dimension *k* to be able to summarize all the drifters in the Arabian Sea region from 2002 to 2017. The predictive dynamics matches the microscopic observations well, and provides us with tremendous confidence to support our macroscopic hypotheses.

Furthermore, the intertwined relationships recovered by the vLDS model between the physical and biological dynamics of the *Noctiluca* blooms, and the intertwined relationships among the biological factors such as *‘chlor_a’, ‘cdm’*, and *‘kd490’* have inspired us to use inference tools to quantify the isolated impact of the biological factors that are responsible for the *Noctiluca* blooms in the Arabian Sea region. The vLDS model presented here is fully generalizable to other datasets for applications in other fields i.e. larval transport etc. in marine larval ecology.

## Acknowledgements

The authors gratefully acknowledge the support from the OceanColor Team, GlobColor Team and Xarray Development Team during the data collection and preprocessing process.

